# Co-translational ribosome pairing enables native assembly of misfolding-prone subunits

**DOI:** 10.1101/2023.06.30.547139

**Authors:** Florian Wruck, Jaro Schmitt, Kai Fenzl, Matilde Bertolini, Alexandros Katranidis, Bernd Bukau, Günter Kramer, Sander Tans

**Author notes:** These authors contributed equally (Jaro Schmitt, Kai Fenzl, Matilde Bertolini).

## Abstract

Protein complexes are pivotal to most cellular processes. Emerging evidence indicates that pairs of ribosomes ubiquitously drive the synchronized synthesis and assembly of two protein subunits into homodimeric complexes^1-5^. These observations suggest protein folding mechanisms of general importance enabled by contacts between nascent chains^6,7^ – which have thus far rather been considered detrimental^8,9^. However, owing to their dynamic and heterogeneous nature, the folding of interacting nascent chains remains unexplored. Here, we show that co-translational ribosome pairing allows their nascent chains to ‘chaperone each other’, thus enabling the formation of coiled-coil homodimers from subunits that misfold individually. We developed an integrated single-molecule fluorescence and force spectroscopy approach to probe the folding and assembly of two nascent chains extending from nearby ribosomes, using the intermediate filament lamin as a model system. Ribosome proximity in early translation stages was found to be critical: when interactions between nascent chains are inhibited or delayed, they become trapped in stable misfolded states that are no longer assembly competent. Conversely, early interactions allow the two nascent chains to nucleate native-like quaternary structures that grow in size and stability as translation advances. We conjecture that protein folding mechanisms enabled by ribosome cooperation are more broadly relevant to intermediate filaments and other protein classes.

## Main

Cells rely on the faithful production of protein complexes. According to textbook models, newly translated polypeptides first undergo a conformational search for the native tertiary structure^10- 12^, and then a diffusion-driven assembly into larger complexes^13,14^. This paradigm is challenged by mounting evidence of co-translational assembly, either between a fully-formed diffusing subunit and a nascent chain^15-20^ (termed “co-post assembly”), or between two nascent chains^1-4^ (termed “co-co assembly”). We recently revealed over 800 co-co assembling homodimers, thus showing the general nature of this biogenesis route^5^. However, the biochemical methods and disome selective ribosome profiling (DiSP)^5^ employed thus far do not detect the nascent chain structures nor the conformational changes during the folding and assembly process. Single-molecule fluorescence and optical-tweezers methods have made important advances in studying nascent chain folding but remain limited to single ribosome-nascent chain complexes (RNCs)^12,21,22^. As a result, we lack insight into the conformational basis and functional relevance of protein complex assembly enabled by coupled ribosomes.

Here we study these issues using the human intermediate filament lamin, whose homodimeric coiled-coil structure represents the largest co-co assembly class^5^. Lamins form a scaffold for the nuclear envelope that spatially organizes chromatin^23,24^, with mutants being the root cause for diseases including premature ageing and cardiomyopathies^25^. Lamin A and its splice variant lamin C contain three domains^26-28^: the N-terminal unstructured head domain, the central ⍰-helical rod domain that mediates dimer formation (Fig. 1A-B, Extended Data Fig. 1), and the C-terminal tail domain with an immunoglobulin-like fold^29^. Lamin polymerization occurs by head-to-tail assembly of lamin homodimers^30^. The lack of lamin heterodimers^31^, even though lamin A and C have identical dimerization domains, led to suggestions of a post-translational sorting mechanism that recognizes the different tail domains^32^. Co-co assembly can alternatively promote isoform-specific homomer formation, and more generally avoid promiscuous interactions between conserved oligomerization domains^33^.

**Fig. 1:**
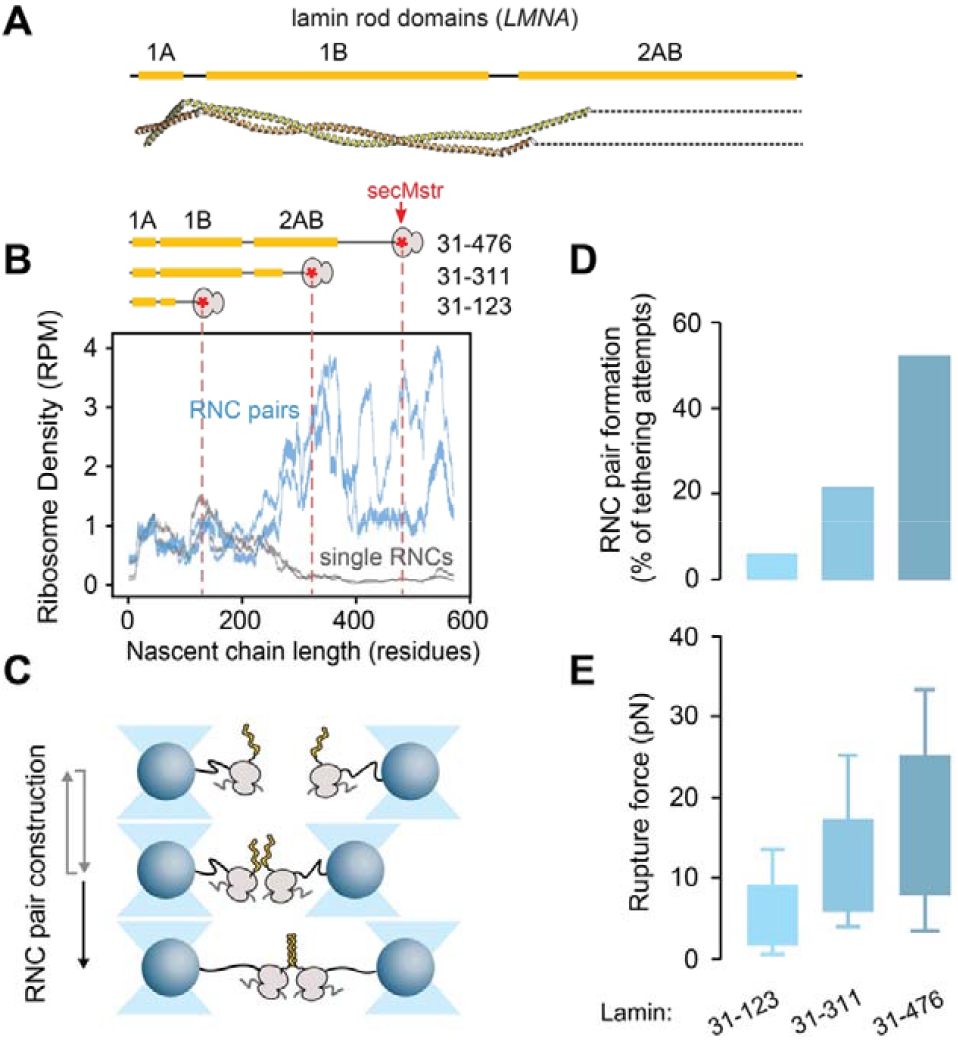
Formation of lamin RNC pairs. **A)** Structure of the lamin A/C homodimer showing the coiled-coil rod domains 1A, 1B and a part of domain 2AB (PDB code: 6JLB^46^). **B)** Lamin RNC pairing *in vivo*. Bottom: Ribosome density along the *LMNA* mRNA for RNC pairs (pairing *via* the nascent chains, blue) and for single RNCs (grey), as obtained by DiSP for U20S cells, with 150 mM KCl and crosslinker present upon cell lysis. Two replicate data sets are shown, as bars representing the position-wise 95% Poisson confidence intervals corrected for library size and smoothed with 15-codon wide sliding window^5^. RPM: reads per million. Top: Lamin nascent chain fragments for optical tweezer experiments (panel C). Dashed lines: ribosome stalling positions. SecMstr: peptide sequence for efficient translation stalling. Yellow bars: coiled-coil domains. Also indicated are the amino acids of the three fragments. **C)** Optical tweezer approach to constitute RNC pairs. Ribosomes, coupled to beads via DNA handles, translate lamin fragments until a secMstr mediated translation arrest using *in vitro* transcription-translation. Next, the beads are repeatedly brought together for 5 seconds, to let nascent chains interact and dimerize, and separated again (grey arrows). A stable tether upon pulling, of twice the DNA handle length, indicates dimer formation (black arrow), while no tether is formed when the chains do not dimerize. **D)** The fraction of dimerization attempts yielding dimer formation, determined as depicted in panel C, for three lamin fragments (n = 136 *LMNA*_31-123_, n = 140 *LMNA*_31-311_, n = 46 *LMNA*_31-476_ cycles). **E)** Rupture force of nascent chain dimer tethers, as measured by ramping up the tether tension, for three lamin fragments (n = 8 *LMNA*_31-123_, n = 30 *LMNA*_31-311_, n = 24 *LMNA*_31-476_ rupturing events).

### *In vivo* detection of lamin nascent chain interactions

We first studied the onset of lamin co-co assembly *in vivo*, including the dependence on cell type (U2OS and HEK cells) and DiSP isolation conditions, which can influence the detected assembly onset (Fig. 1B, Extended Data Fig. 2). The DiSP method is based on isolating RNC pairs that are coupled via their nascent chains (disomes), as well as uncoupled RNCs (monosomes), followed by sequencing their protected RNA footprints as in standard ribosome profiling. Consistently, for lamin RNC pairs, the number of mRNA reads increased after synthesis of coil 1B (after 250 residues), as the reads decreased for the uncoupled RNCs^5^. The increased level of RNC pairs persisted during synthesis of the rod domain, while the uncoupled RNC level did not recover. This transition was not affected by salt or crosslinking conditions (Extended Data Fig. 2), indicating robustness of lamin assembly onset detection using DiSP. However, such approaches to purify RNC pairs from cells are less suited to study nascent chain conformations^5,34,35^, due to RNC heterogeneity and difficulty of incorporating measurement probes. Hence, we aimed to construct RNC pairs *in vitro*, stalled at key phases of translation identified by the DiSP data: before the onset of dimerization, with only the small ⍰-helical coil 1A fully translated, after the dimerization onset with coils 1A and 1B fully translated, and finally after translation of coil 2AB, with the full-length rod domain translated and ribosome exposed (Fig. 1B).

**Fig. 2:**
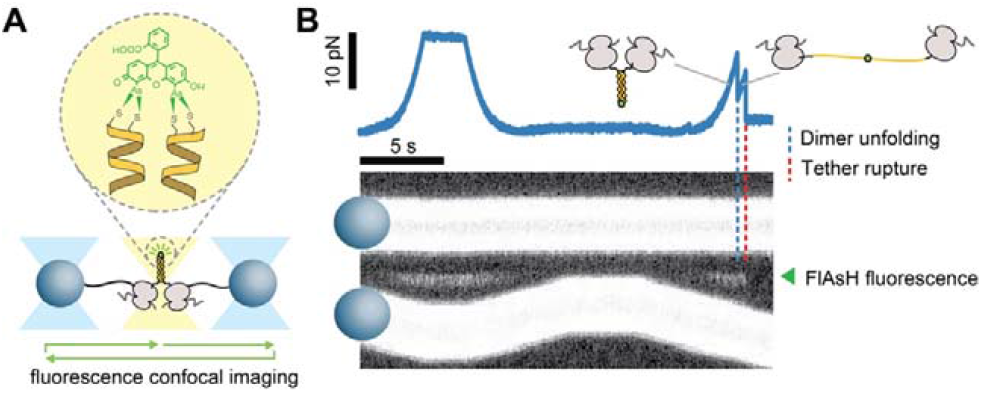
Fluorescent detection of nascent lamin dimers. **A)** Approach for visual verification of N-terminal co-localization. Two nascent chain N-termini can be linked by the FlAsH dye, if they co-localize as in the native coiled-coil dimer structure (Fig. 1A), using two N-terminally incorporated cysteines. Bound fluorescent FlAsH is detected by scanning a confocal excitation beam (yellow beam, green arrows) along the molecular tether, while the optical tweezers laser beams (blue) trap the beads. **B)** Corresponding data. Bottom: detected fluorescence scans in time, showing bead movements in stretch-relax cycles and the FlAsH fluorescence signal between the two beads (green arrowhead), when the tether is under tension and hence stably in focus. RNC pair construct: *LMNA*_31-476_. Top: corresponding measured force acting on the beads and tether. Blue dashed line: sudden drop in force indicates unfolding, while FlAsH keeps the N-termini connected (see panel A). Red dashed line: tether rupture, likely by dissociation of one of the DNA handles from the bead surface.

### *In vitro* formation of lamin RNC pairs

To construct pairs of RNCs coupled by their nascent chains, we first linked biotinylated ribosomes to polystyrene beads via 5 kbp DNA handles (Fig. 1C). Synthesis of lamin nascent chains by the bead-tethered ribosomes was performed by *in vitro* transcription-translation, using the ‘SecM strong’ sequence to stall translation at positions indicated above (Fig. 1B, Extended Data Fig. 3). Two such beads were captured by two optical traps, repeatedly brought together, within about 200 nm for about 5 seconds, and separated again. We quantified the fraction of approach-retract cycles in which a tether formed between the beads that was twice the length of a single DNA handle (Extended Data Fig. 4), and hence indicated coupling of two RNCs via their nascent chains (Fig. 1C). This fraction increased from below 10% to above 50% with increasing fragment length (Fig. 1D). Such an increasing trend agrees with the lamin DiSP data (Fig. 1B), and with coupling taking place *via* the nascent chains, rather than *via* other (ribosomal) components. While rare, we did detect dimerization events already for the shortest fragment, indicating an earlier assembly onset than detectable by DiSP (Fig. 1B). Shorter nascent chain dimers may be less stable and therefore escape detection by DiSP. To probe the stability of the nascent chain dimers against dissociation, we measured the force required to rupture the tether by increasing the distance between the beads. The tethers ruptured in a single step, in line with coiled-coils unfolding and dissociating discretely during pulling^36^. The rupture force indeed increased with fragment length, from about 5 pN to over 15 pN (Fig. 1E), consistent with a progressively larger coiled-coil dimer interface. Overall, the data indicated that lamin nascent chains dimerized when brought in close proximity, with loose associations starting at short chain lengths and increased stability with increasing chain length.

**Fig. 3:**
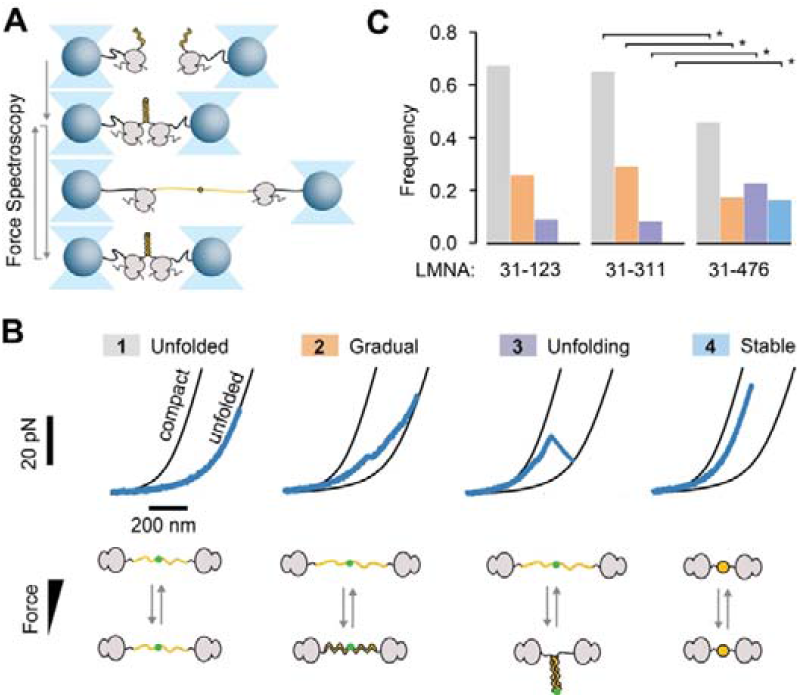
Delayed RNC interaction promotes non-native lamin dimer conformations. **A)** Diagram of nascent chain dimer force spectroscopy approach. Lamin nascent chain dimers are formed as depicted in Fig. 1 C, and subsequently exposed to repeated stretching and relaxation while the split-FlAsH-tag (green dot, see also Fig. 2A) keeps N-termini bound together upon unfolding. **B)** Classes of force-extension behaviour of lamin nascent chain dimers. Black lines are theoretical worm-like chain (WLC) curves for both nascent chains being compact (left) or fully unfolded (right). The position of the measured data (blue) in between these two reference curves indicates what fraction of the nascent chain (how many amino acids) is in the compact state, and what fraction is in the extended state. During stretch-relax cycles four classes are observed, with the majority of the chain: (1) remaining unfolded, (2) initially compact and extending and compact gradually under tension, as expected for linear 17-helices, (3) initially compact and unfolding discretely below 45 pN, as expected for coiled-coil dimer structures, (4) initially compact and remaining so up to 45 pN, typically for multiple stretch-relax cycles, indicative a kinetically trapped misfolded state (3). Data is shown for *LMNA*_31-476_ fragments. **C)** Frequency of stretch-relax cycles with observed force-extension features (see panel B), for lamin nascent chain dimers of three nascent chain lengths (n = 14 *LMNA*_31-123_ molecules, 24 pulling cycles, n = 38 *LMNA*_31-311_ molecules, 53 pulling cycles, n = 65 *LMNA*_31-476_ molecules, 112 pulling cycles). Dimer formation and misfolding increases in frequency with increasing nascent chain length, at the expense of unfolded states. Star: significant difference (p < 0.05). Colours are as in panel B.

**Fig. 4:**
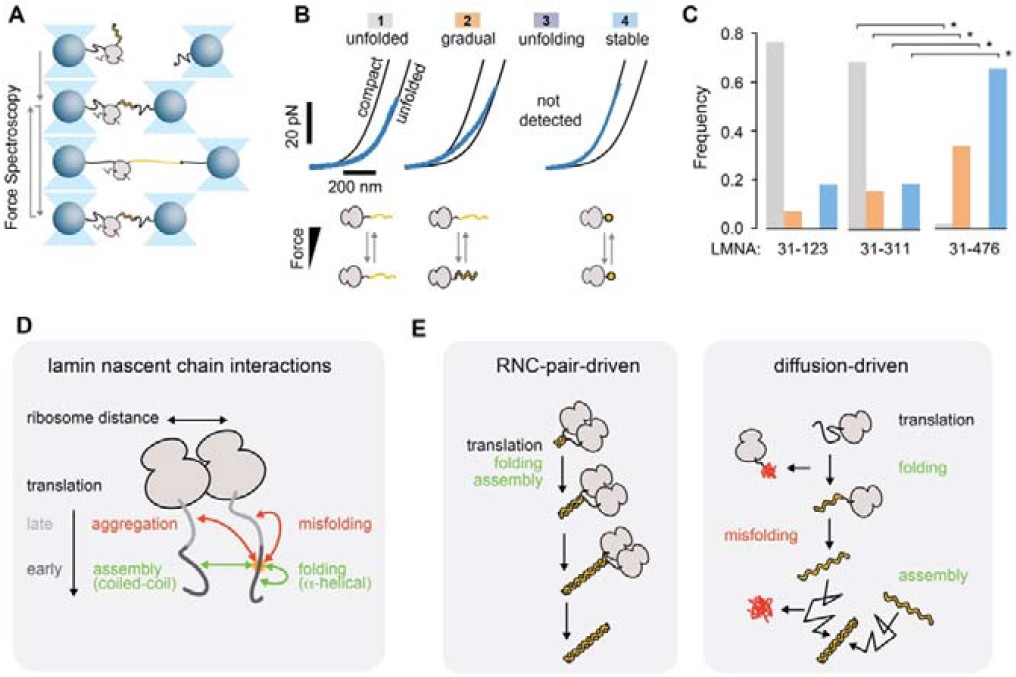
Lamin misfolds when interactions with other nascent chains are denied. **A)** Diagram of optical tweezer approach to probe single monomeric lamin nascent chains. Stalled ribosomes, coupled to beads via DNA handles, translate lamin nascent chain fragments, incorporating biotin N-terminally. This biotin is coupled to a second trapped bead via another DNA handle. The nascent chain conformations are hence probed by repeated stretching and relaxation (grey arrows). Indicated is the transition between the unfolded and linear ⍰-helical conformations. **B)** Classes of observed force-extension behaviour for monomeric lamin nascent chains. Black lines indicate reference behavior for a single nascent chain being compact (left) or fully unfolded (right). During stretch-relax cycles four classes of conformational states are distinguished, with the majority of the chains: (1) remaining unfolded, (2) being initially compact and extending gradually under tension, as expected for linear ⍰-helices, (3) being initially compact and unfolding discretely below 45 pN, as expected for coiled-coil dimer structures, (4) being initially compact and remaining so up to 45 pN, typically for multiple stretch-relax cycles, indicative of a kinetically trapped misfolded state. Data is shown for *LMNA*_31-476_ fragments. **C)** Frequency of stretch-relax cycles with specific force-extension features (see panel B), for lamin nascent chain monomers of three nascent chain lengths (n = 14 *LMNA*_31-123_ molecules, 45 pulling cycles, n = 13 *LMNA*_31-311_ molecules, 32 pulling cycles, n = 19 *LMNA*_31-476_ molecules, 67 pulling cycles). Star: significant difference (p < 0.05). **D)** Competition between initial lamin nascent chain interactions and factors that affect it. A given site (orange) can engage in non-native aggregation and misfolding interactions (orange arrows), and native (⍰-helical) folding and (coiled-coil) assembly contacts (green arrows). The two lamin coils natively contact ‘in-register’, starting with the co-localizing N-termini (Fig. 1A) at an early time point of translation. Delay of inter-chain interactions until later phases of translation results in more competition from ‘out-of-register’ aggregation and misfolding interactions between earlier and later translated segments. Larger ribosome distances limit inter-chain interactions, which inhibits the stabilizing assembly interactions, and hence promotes intra-chain misfolding interactions. **E)** Schematic representation of RNC-pair- and diffusion-driven complex formation. Two nascent chains may either be synthesized first and then dimerize by diffusion (right), or dimerize during translation by two proximal ribosomes (*co-co* assembly, left). The latter promotes proper folding and dimer assembly, by establishing native inter-chain contacts before non-native intra-chain contacts can form, and by limiting cytosolic exposure of protein chain monomers during translation and diffusion. Co-co assembly of lamin coiled-coils results in progressive dimer growth, in a process that merges ⍰-helical folding and assembly and is driven by ongoing translation.

### Nascent lamin complex formation observed by fluorescence

To test whether the observed nascent dimers are consistent with the native coiled-coil lamin structure, we integrated fluorescence detection into the optical tweezers assay (Fig. 2A). We inserted a pair of adjacent cysteines at the lamin N-terminus and reasoned that two such pairs should co-localize in the native parallel coiled-coil conformation. Hence, a bipartite tetra-cysteine motif would form that can bind the FlAsH dye^37^, which becomes fluorescent upon binding, and detectable by confocal fluorescence imaging. We performed the above approach-retraction protocol to form a lamin RNC pair and exposed it to FlAsH in solution. To detect bound FlAsH, a confocal fluorescence excitation beam was scanned across the beads and along the connecting tether. A fluorescent signal indeed appeared in-between the beads, visible as an additional line in kymographs that display the subsequent fluorescence emission scans (Fig. 2B). The two lamin N-termini were thus in close proximity, in line with the native coiled-coil structure of lamin dimers (Fig. 1A). The longer fragments again showed higher dimer formation frequency (Extended Data Fig. 5), consistent with the previous measurements in the absence of FlAsH (Fig. 1D).

### Delayed RNC interaction promotes non-native lamin dimer conformations

The above experiments showed that dimers of nascent lamin chains can be formed *in situ* but did not provide insight into unsuccessful assembly attempts or folding errors. We surmised the split FlAsH-tags may keep the N-termini connected after full unfolding, which could allow us to directly follow dimer formation and dissociation, including underlying conformational changes. Upon RNC coupling and FlAsH exposure, tethers indeed remained intact after lamin dimers were unfolded by stretching, as evidenced by a force drop to a non-zero level and continued presence of the fluorescence signal (Fig. 2B). Note that the fluorescence signal is not visible at the lowest forces because the molecular tether is then not properly positioned in the confocal imaging plane. The force and fluorescence signals disappeared when the tether fully ruptured, which could be due to FlAsH, Dig-AntiDig, or biotin-neutravidin dissociation, but whose rupture force should not depend on nascent chain length. Accordingly, we found similar rupture forces for all three constructs (Extended Data Fig. 6). In this manner, nascent chain dimers could be cyclically unfolded by stretching and then given a chance to reform by relaxation (Fig. 3A).

Four classes of behaviour were observed (Fig. 3B, Extended Data Fig. 7), with individual RNC pairs showing different classes from one cycle to the next (Extended Data Fig. 8). The chains were for the largest part either: (1) initially unfolded, and remaining so throughout the stretch-relax cycle, (2) initially compact and gradually decompacting during stretching (or conversely becoming gradually more compact during relaxation), with the data resembling previous pulling experiments on stretched linear ⍰-helices^38,39^, (3) initially compact and unfolding discretely, consistent with the formation and unfolding of the coiled-coil dimer^40^ (see also Fig. 2B), or (4) initially compact and remaining so throughout the cycle, with the applied force unable to unfold the structure.

The compacted states (4) indicated the formation of stable non-native conformations that differ from coiled-coil or linear ⍰-helical structures^41^. These states were preserved for multiple cycles until the tether ruptured, indicating that the nascent chains could no longer form coiled-coil-like dimers. Conversely, unfolded (1) and ⍰-helical-like states (2) could transition to the coiled-coil-like state that unfolded in discrete steps (3) (Extended Data Fig. 8). In line with these data and the increased dimerization propensity with nascent chain length (Fig. 1D), class (3) was found to increase in frequency with nascent chain length, while classes (1) and (2) decreased (Fig. 3C). Notably, class (4) was only observed for the longest fragment (Fig. 3B, C, Extended Data Fig. 7). Note that the two nascent chains are in late stages of translation when they interact in these relax-stretch experiments. Overall, these data indicated that two neighbouring nascent chains can adopt ⍰-helical and coiled-coil dimer structures early during translation, which grow as translation progresses. Alternatively however, non-native states are promoted when the interactions between chains are delayed until later phases of translation.

### Co-co assembly supresses intra-chain lamin misfolding

A key question concerns the conformational competition that determines the different observed folding pathways. Native complex formation can be in competition with aggregation interactions between the chains, or misfolding interactions within chains – both of which could yield the observed non-native states (Fig. 3). To address this issue, we probed single ribosome-associated lamin nascent chains, in absence of partnering RNCs (Fig. 4A). First, we generated stalled ribosomes tethered to beads via 5 kbp DNA handles as before, while using suppressor tRNAs to biotinylate the nascent chains N-terminally. After trapping one bead, the biotinylated nascent chain was linked to a second trapped bead via another 5 kbp DNA handle. Next, we cyclically first separated and subsequently approached the two optical traps, thus stretching and relaxing the ⍰-helical lamin rod domains by their N-and C-termini, for all three lamin fragments (Fig. 4A).

With single RNCs, we observed three of the four classes (Fig. 4B-C, Extended Data Fig. 9): (1) unfolded, (2) ⍰-helix-like, and (4) non-native. As expected, the discrete unfolding of the coiled-coil-like dimer class (3) was not detected. Most notable was the prominence of the non-native class (4). This class was observed for all fragments, at frequencies ranging from 20% for the shortest to 60% for the longest (Fig. 4C), while for the RNC pairs it was detected for the longest fragment only, and at a low frequency of 20% (Fig. 3C). These data indicate that in absence of another lamin nascent chain, the individual lamin nascent chains were prone to form compact and stable non-native structures. Indeed, ⍰-helices can form misfolds that have high contact-order and are hence compact^42^. Conversely, the presence of a second nascent chain suppressed this lamin misfolding while enabling the coiled-coil-like assembly. Hence, native structure formation here is not inhibited but instead promoted by interactions between nascent polypeptide chains.

## Discussion

In this study, we report that by coupling co-translationally, RNC pairs can drive the assembly of coiled-coil homodimers composed of subunits that misfold individually (Fig. 4D). We find that RNC proximity and translation biases the competition between native and non-native contacts, which both can form either within single chains or between two chains. Specifically, residues that are translated early can form non-native contacts with chain segments that are translated later (Fig. 4D, red arrows), which compete with native coiled-coil contacts (Fig. 4D, green arrows). Timely formation of the latter native contacts between nascent chains allows the realization of partial coiled coils, which grow in size and stability as translation proceeds (Fig. 4E). Conversely, non-native contacts are promoted when the RNCs either cannot interact, or interaction is delayed until later phases of translation. Lamin nascent chains then misfold and are no longer assembly competent (Fig. 4B-C). These risks should be further aggravated when assembly is postponed until after translation, and the monomeric lamin subunits must diffuse through the cytosol in order to dimerize (Fig. 4F). Overall, nascent chain interactions during translation thus provide a reciprocal chaperoning function that suppresses misfolding, in contrast with the view that high local nascent chains concentrations promote aggregation^43^.

The ability to synthesise complexes composed of misfolding-prone subunits may be of broad relevance to cells, as it expands the range of possible protein structures and functions. Indeed, the elongated lamin shape renders the lamin subunits prone to misfolding in monomeric form, yet is also key to its unique mechanical properties^44^. The underlying mechanism of coupled folding and assembly by RNC pairs is likely more widely relevant, given the recently observed prevalence of RNC pairing for coiled-coil and other homodimer classes, including BTB and Rel homology domain proteins^5^, and may extend to heterodimers, higher-order oligomers, as well as membrane-based biogenesis. Hence, it may contribute to understanding the architecture and (mutation-induced) misfolding of proteins ranging from intermediate filaments to major regulatory and metabolic proteins such as initiation factor 2B and Fatty Acid Synthase. Furthermore, the findings raise the question whether the formation and translation synchrony of RNC pairs is controlled. The latter may be achieved by sequence-induced translational pausing or regulatory factors, as recently studied in ribosome collision detection^45^. Finally, the results may open up new routes in artificial protein and mRNA therapy design.

## Acknowledgements

Work in the group of S.J.T. is supported by the Netherlands Organization for Scientific Research (NWO). M.B. and K.F. were supported by a HBIGS PhD fellowship. M.B. was additionally supported by a Boehringer Ingelheim Fonds (BIF) PhD fellowship. F.W. received funding from the European Union’s Horizon 2020 Research and Innovation Programme under the Marie Sklodowska-Curie grant agreement No 745798. This work was supported by the Helmholtz-Gemeinschaft (DKFZ NCT3.0 Integrative Project in Cancer Research (DysregPT_Bukau 1030000008 G783)), the European Research Council (ERC Advanced grant (743118), and the Klaus Tschira Foundation. J.S., K.F. and M.B. are members of the Heidelberg Biosciences International Graduate School (HBIGS).

## Author Contributions

Conceptualization: F.W., J.S., K.F., M.B., B.B., G.K., S.J.T. Methodology: F.W., J.S., K.F., M.B., B.B., G.K., S.J.T. Biotinylated ribosomes: A. K. Experiments: F.W., J.S. Formal analysis, and data visualization: F.W., K.F., M.B., B.B., G.K., S.J.T. Writing: all authors. Supervision: S.T., B.B. and G.K.

## Declaration of Interests

The authors declare no competing financial interests. Correspondence and requests for materials should be addressed to S.J.T.

## Notes

### Competing Interest Statement

The authors have declared no competing interest.

