## Supplemental information for "Co-translational ribosome pairing enables native assembly of misfolding-prone subunits"

**Extended Data Figures**

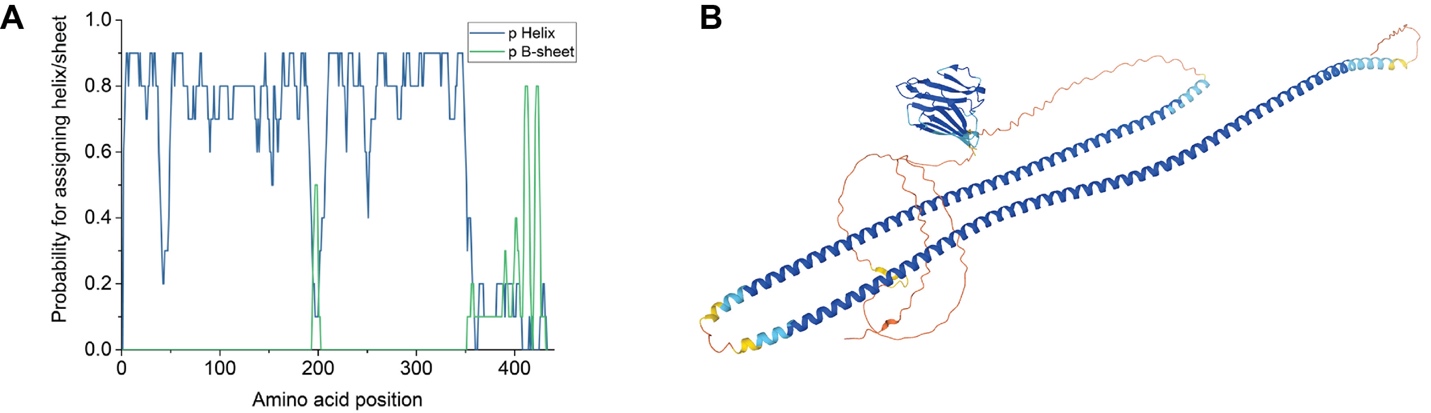

**Extended Data Fig. 1. Structure prediction of Lamin. A)** Estimation of the secondary structure of *LMNA*_31-476_ using PredictProtein^1^. The N-terminal ~350 amino acids of this protein show a high probability to form an alpha helix, as expected from the known coiled-coil quaternary structure of the lamin rod domain in lamin dimers (Fig. 1A). **B)** Structure prediction of Lamin A by AlphaFold (Uniprot P02545), indicating the alpha-helical rod domain, the C-terminal domain with immunoglobulin-like fold and unstructured segments. Colors indicate model confidence, from dark blue (very high) to orange (very low).

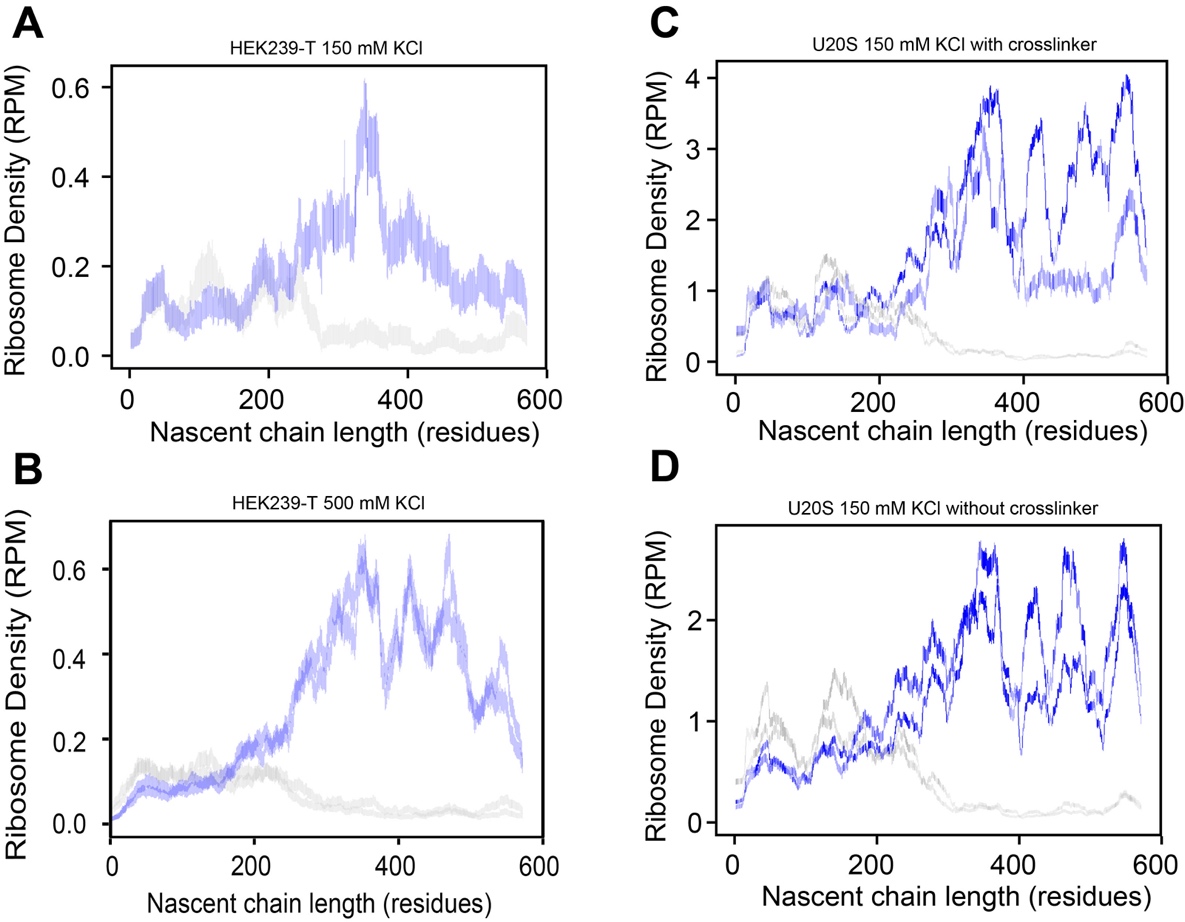

**Extended Data Fig. 2. *In vivo* detection of Lamin nascent chain interactions.** Lamin DiSP curves (codons 1-572) from different cell types, crosslinking, and salt conditions. Blue: density of ribosomes engaged in RNC pairs (coupled by their nascent chains) positioned along the *LMNA* mRNA (quantified as lamin nascent chain length). Gray: density of single ribosomes (not in pairs). Data are plotted as vertical bars, indicating the position-wise 95% Poisson confidence intervals corrected for library size and smoothed with a 15-codon wide sliding window. The color intensity of the bars represents the amount of reads per position^2^. **A)** Lamin data from HEK 239-T cells, with 150 mM KCl during lysis (two replicates). **B)** Lamin data from HEK 239-T cells, with 500 mM KCl during lysis (two replicates). Data reproduced with permission^2^. **C)** Lamin data from U20S cells with 150 mM KCl during lysis and crosslinked nascent chains (two replicates, crosslinked with 2.5 mM BS3 and 20 mM EDC). **D)** DiSP data from U20S cells with 150 mM KCl during lysis, without crosslinkers (two replicates). Overall, the curves show that the onset of assembly is similar for all data sets.

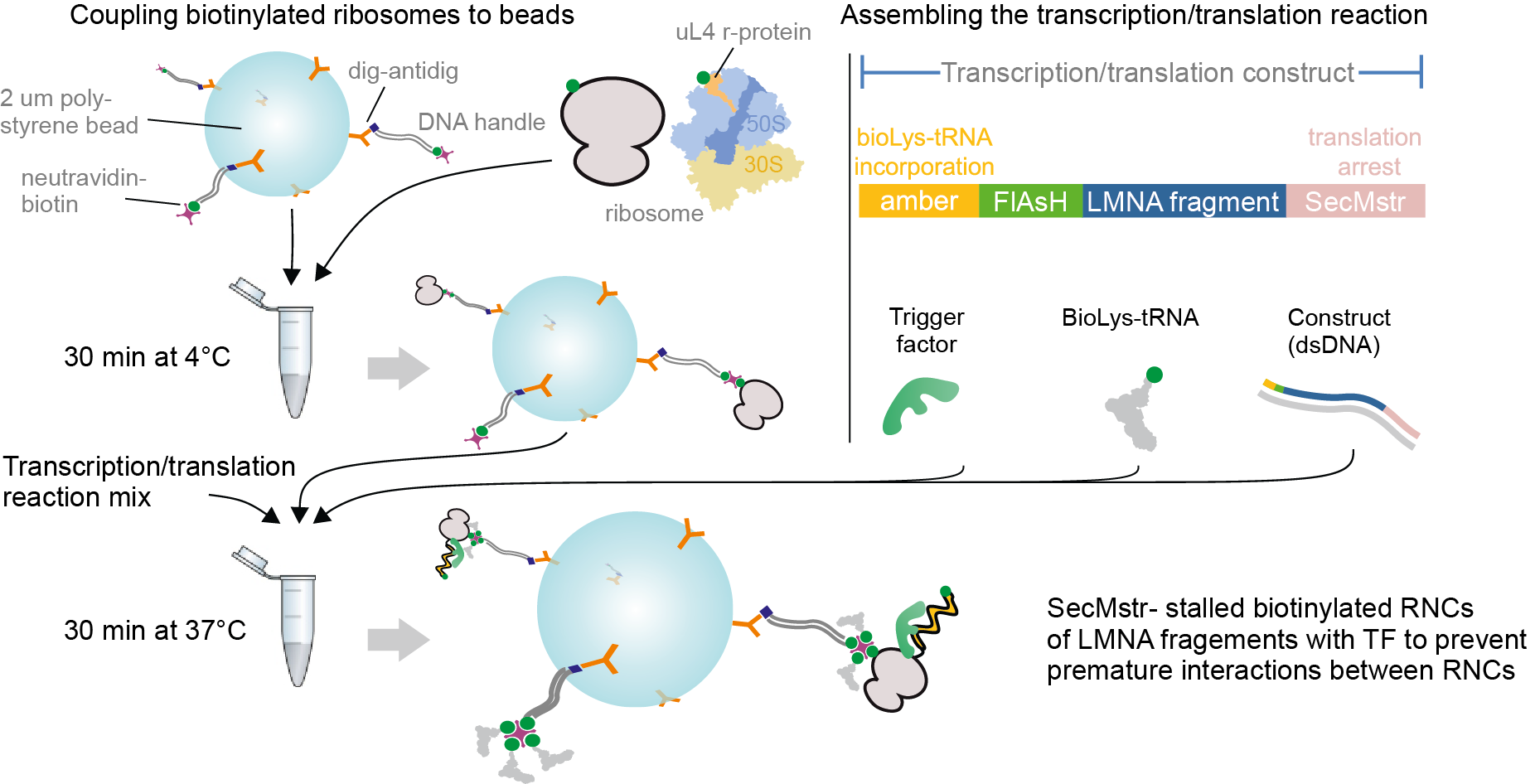

**Extended Data Fig. 3. Schematic of the *in-vitro* RNC generation protocol.** First, biotinylated ribosomes are coupled to Neutravidin-DNA (5 kbp long) coated beads. Next, SecMstr-stalled nascent chains of either *LMNA*_31-123_, *LMNA*_31-311_, or *LMNA*_31-476_ fragments are synthesized using a customized PURE expression system, which included the chaperone trigger factor to mitigate bead clustering via nascent-chain interactions. Forming a pair of two ribosome nascent chain complexes (RNCs) is described in the main text and methods. Briefly, two such beads are brought together such that the RNCs can interact, and then retracted to assess whether a tether of twice the size of a DNA handle is formed (Extended Data Fig. 4). When the two corresponding nascent chains form a coiled-coil structure, their N-termini co-localize. These N-termini each contain two adjacent cysteines, such that a bipartite tetra-cysteine motif forms that binds a FlAsH dye^3^. In order to probe RNC monomers, their nascent chain N-terminus is linked to a DNA handle. For this purpose, biotin is incorporated co-translationally at the N-terminal amber position TAG using a modified tRNA pre-charged with biotinylated lysine, which is linked to the DNA handle using neutravidin. See methods for further details.

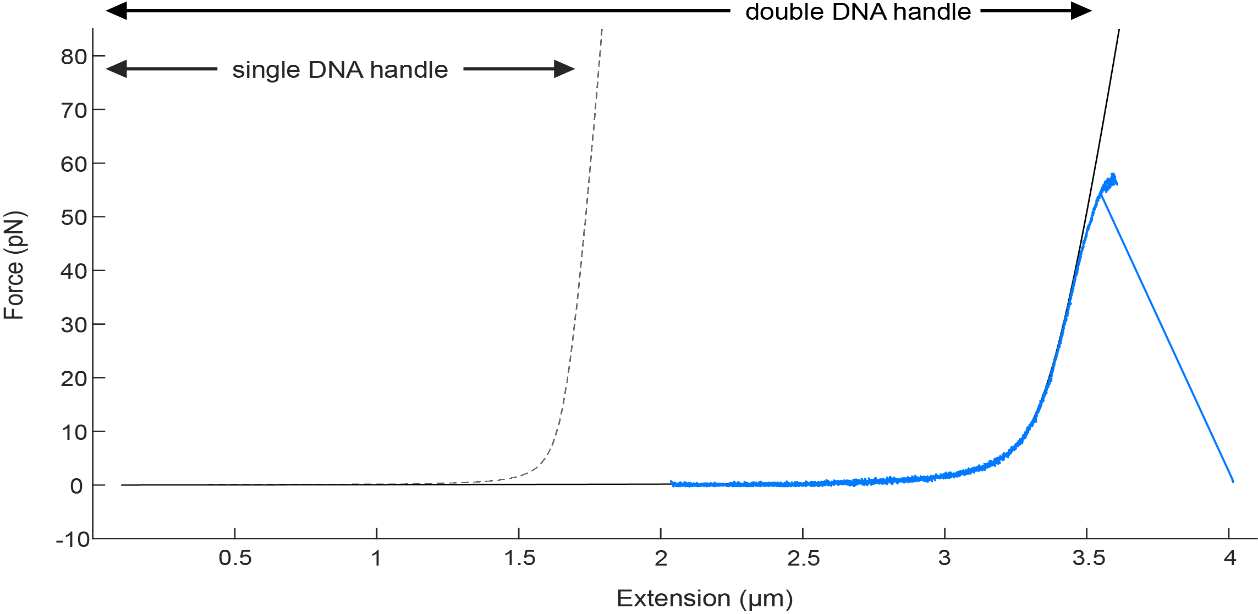

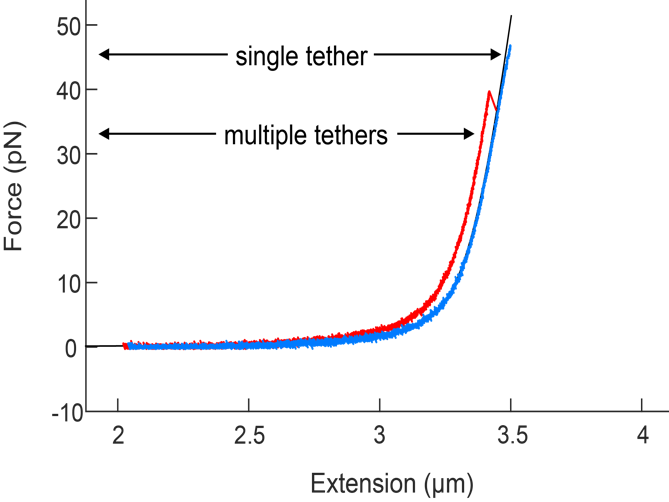

**Extended Data Fig. 4. Establishing single tethered RNC pairs.** Top: The DNA handles used for tethering the RNCs have a length of 5 kbp (dashed black curve, eWLC model for 1.7 µm contour length DNA). To form an RNC pair, two beads coated with DNA-tethered single RNCs are brought together such that the nascent chains can interact. A tether is then formed (blue curve) with a length corresponding to two DNA handles (3.4 µm contour length), which exhibits Force-Extension characteristics (blue) as predicted by the eWLC model (solid black curve) within a force range of 0-35 pN^4^. Above ~35 pN individually tethered molecules begin to deviate slightly from the eWLC model due to twisting of the DNA handles. Bottom: More than one such tethers can form between the optically trapped beads, which are not aligned in parallel with the bead-bead axis resulting in a shorter bead-bead distance, do not follow the eWLC model, and do not yield a single step break when one tether breaks (red), and can hence be identified.

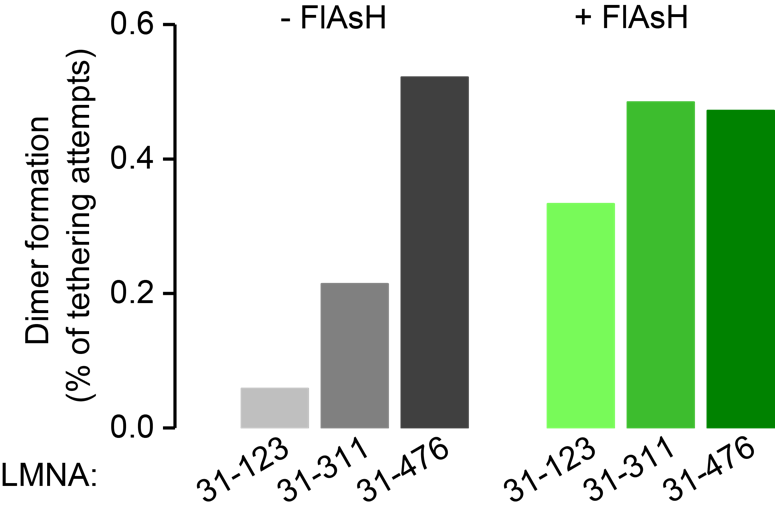

**Extended Data Fig. 5. RNC dimer formation frequency.** Dimer formation as a fraction of dimerization attempts in the presence (n = 91 molecules) or absence (n = 62 molecules) of FlAsH for all lamin C nascent chain fragments. Data in absence of FlAsH (grey), also plotted in Fig. 1D, is added for comparison.

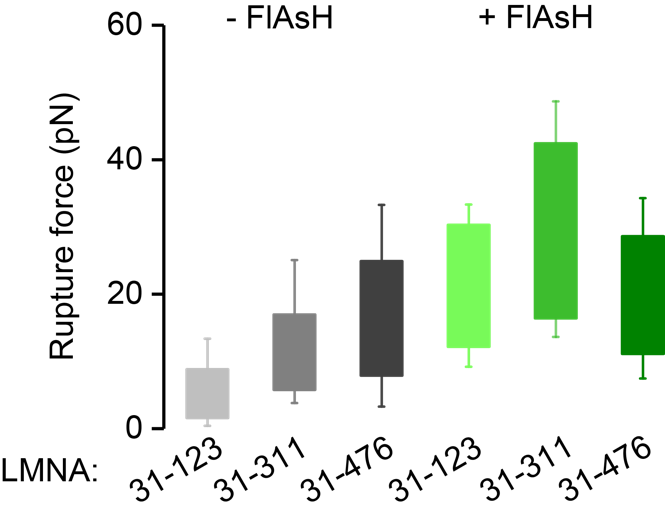

**Extended Data Fig. 6. RNC dimer rupture force during stretching.** Dimer rupture forces in the presence (n = 91 molecules) or absence (n = 62 molecules) of FlAsH for all lamin C nascent chain fragments. Data in absence of FlAsH (grey), also plotted in Fig. 1E, is added for comparison.

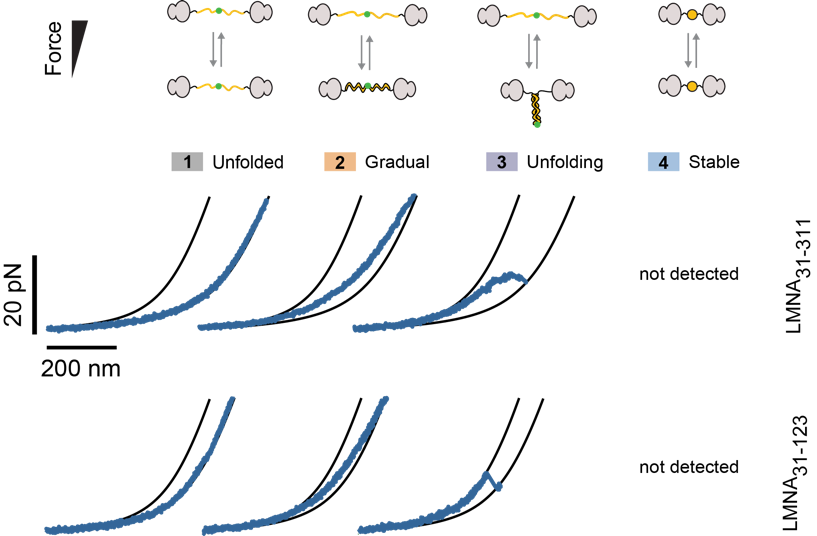

**Extended Data Fig. 7. Classes of force-extension behavior of the lamin nascent chain dimers *LMNA*_31-311_ and *LMNA*_31-123_**. Black lines are theoretical worm-like chain (WLC) curves for both nascent chains being compact (left) or fully unfolded (right). The position of the measured data (blue) in between these two reference curves indicates what fraction of the nascent chain (how many amino acids) is in the compact state, and what fraction is in the extended state. During stretch-relax cycles of these two lamin fragment lengths three classes are observed, with the majority of the chain: (1) remaining unfolded, (2) initially compact and extending and compact gradually under tension, as expected for linear ⍺-helices, and (3) initially compact and unfolding discretely below 45 pN, as expected for coiled-coil dimer structures. The stably compacted class (4), detected during measurements of the longest lamin fragment *LMNA*_31-476_ (Fig. 3B), was not observed here, consistent with misfolding being suppressed by nascent chain interactions.

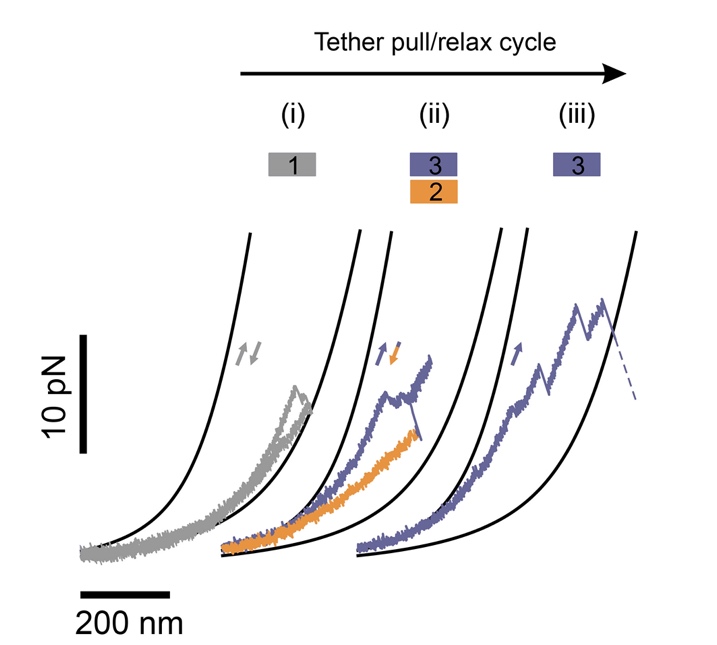

**Extended Data Figure 8. Sequence of conformational transitions.** Three consecutive pull-and-relax cycles of a RNC dimer (*LMNA*3_1-476_), showing subsequent conformational transitions, which have a heterogeneous character at the molecular level. Transitions corresponding to three different folding features (1-3) are observed during the three cycles (i-iii). In the first pull the dimer is mostly unfolded (gray), then folds after relaxation to 0 pN, as observed in the second pulling cycle, where it unfolds partially during pulling and relaxation (purple), before gradually compacting during relaxation (orange). In the third cycle the molecule is found to be compact, and then unfolds once more before rupturing during stretching (purple).

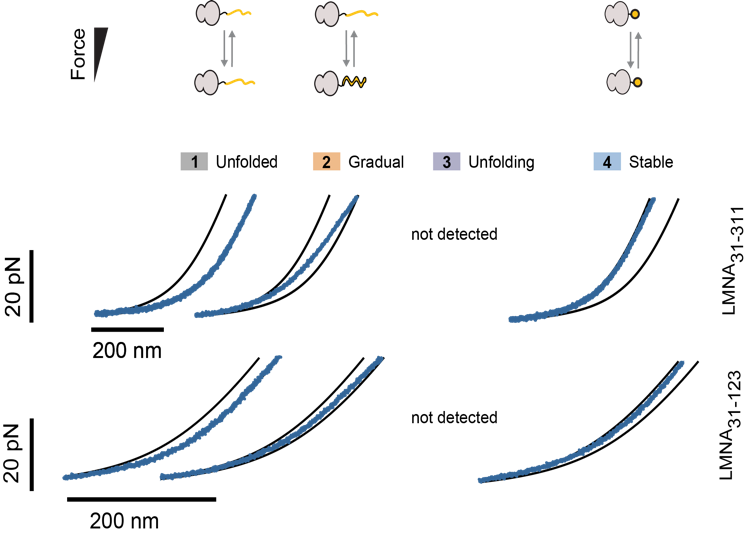

**Extended Data Fig. 9. Classes of force-extension behavior of lamin nascent chain monomers *LMNA*_31-311_ and *LMNA*_31-123_.** Black lines are WLC reference curves for the single nascent chain being compact (left) or fully unfolded (right). During stretch-relax cycles four classes of conformational states are distinguished, with the majority of the chains: (1) remaining unfolded, (2) being initially compact and extending and compact gradually under tension, as expected for linear ⍺-helices, (3) being initially compact and unfolding discretely below 45 pN, as expected for coiled-coil dimer structures, (4) being initially compact and remaining so up to 45 pN, typically for multiple stretch-relax cycles, indicative of entry into a kinetically trapped misfolded state (3). Class 3 is not observed for any fragment, which is consistent because the coiled-coil cannot form without the second chain present.

**METHODS**

**Disome Selective Profiling (DiSP)**

U2OS (ATCC Cat# HTB-96, RRID: CVCL_0042) and HEK293-T cells (DSMZ Cat# ACC 635) were cultivated in high glucose DMEM media containing GlutaMAXTM and pyruvate (Gibco), which was freshly supplemented with 10% heat-inactivated FCS (Gibco), 100 units/mL penicillin and 100 μg/mL streptomycin (Gibco) and were grown in a humidified incubator with 5% CO2 at 37°C (HERAcell 150i). Variations in the lysis protocol were implemented in different datasets. A lysis buffer with physiological salt concentration (50 mM HEPES pH 7.0, 10 mM MgCl_2_, 150 mM KCl, 1% NP40, 10 mM DTT, 100 μg/ml CHX, 25 U/ml recombinant Dnase1 (Roche) and protease inhibitor (complete EDTA free, Roche)) was used for DiSP of HEK293-T and U2OS cells (Fig. S2 A and C). These DiSP gene density profiles show the position-wise 95% Poisson confidence interval corrected for library size, and read counts are smoothed with a 15-codon wide sliding window^1^. A high-salt lysis buffer containing 500 mM KCl was employed for DiSP of HEK293-T cells to test possible effects on onset of lamin disome formation (Fig. 1B). DiSP of U2OS cells was also performed by lysing cells in the presence of chemical crosslinkers (Fig. S2 B). In this case, the lysis buffer was additionally supplemented with 2.5 mM BS3 and 20 mM EDC. Cells were scraped in cross linkers-containing lysis buffer on ice, so that crosslinking occurred simultaneously to lysis.

**Cloning**

All primer sequences used for cloning are available in Table S1. *LMNA* corresponding to lamin C that lacks the unstructured head domain (residues 31-542), was PCR-amplified from a self-made U2OS cDNA library (SuperScript™ III first-strand synthesis kit, ThermoFisher). The employed PCR primers (MB143 + MB144) added an NdeI restriction site followed by a splitFlAsH tag (SF: MAGSCCGG) at the 5’ end and a TwinStrep tag (TS: GGSGSAWSHPQFEKGGGSGGGSGGSAWSHPQFEKGA) with a BamHI overhang at the 3’ end of the construct (final sequence named SFLMNCTS available in Table S2). T4 DNA ligase was used to ligate the gel-purified PCR fragment into a BamHI/NdeI restricted pET3a vector. The resulting plasmid was sequenced with standard Eurofins primers (T7 forward and pET reverse primers) and custom primers (MB75 + MB76). Fragments for Gibson assembly were generated by PCR reactions on the SF-LMNC-TS amplicon (described above) using the primer combinations JS9 and JS10, JS9 and JS11, JS9 and JS12. Plasmid pRSET-AceE(1-181) served as a template in a PCR reaction with primers JS7 and JS8 to create the linear plasmid backbone, containing the SecMstrong sequence (FSTPVWIWWWPRIRGPP) at the 5’ end and the T7 promoter at the 3’ end. The *LMNA* containing fragments were fused to the linear plasmid backbone by Gibson assembly. 5 μL of the reaction mix was used for transformation into 5-alpha competent *E. coli* (NEB) according to the manufacturers' protocol and selected on LB plates containing 100 μg/mL ampicillin. Plasmids were isolated using the QIAprep Spin Miniprep Kit (QIAGEN). The templates for the in vitro transcription/translation reaction were amplified by PCR from the generated plasmids using primers JS28 and JS29.

Coupling of neutravidin-DNA handles to beads

Double-stranded DNA (dsDNA) molecules were prepared by PCR amplification using digoxigenin (DIG) and biotin 5’-end-modified primers. 2DIGfw5kbp and 3BIOrev5kbp were used with pOSIP-TT as a template in a two-step Phire Green Hot Start II PCR (Thermo Scientific) reaction. The 5 kb PCR product was purified using the QIAquick PCR Purification Kit (QIAGEN). 10 nM 5 kb DNA was coupled to 2 μM neutravidin (Thermo Scientific) by incubation in 10 mM PBS overnight at 4°C. 2.1 μm diameter carboxyl-functionalized polystyrene beads (Spherotech) were modified with anti-digoxigenin (anti-DIG, Roche), using the carbodiimide crosslinker EDAC, following the PolyLink protein coupling kit protocol (Polysciences). Subsequently, the resulting 5 kbp neutravidin-DNA handles (1.7 µm contour length) were coupled to the anti-DIG beads at a reaction ratio of ~10 neutravidin-DNA/bead for 30 min at 4°C. The beads with the neutravidin-DNA handles were then washed several times with Tico buffer (20 mM HEPES-KOH pH 7.6, 10 mM (Ac)_2_Mg, 30 mM AcNH_4_, 4 mM β-mercaptoethanol) and split into two batches (modified protocol from^2^).

Coupling of ribosomes to beads with DNA handles:

Ribosomes from an RNase deficient *E. coli* K-12 strain (Can20/12E^3^) were biotinylated *in vivo* at the uL4 ribosomal protein and subsequently isolated^4^. Biotinylated ribosomes were added to one batch of Tico-washed neutravidin-DNA modified beads at 350 nM, supplemented with murine RNase inhibitor (New England Biolabs) and incubated at 4°C for 30 min. The remaining unbound ribosomes were then removed via pelleting and the beads were washed once with Tico buffer before they were resuspended directly into the cell-free transcription/translation mix described below.

Cell-free protein synthesis and co-translational labelling

The cell-free transcription/translation mix used in this study is a customized version of the bacterial PURE system without ribosomes (PUREexpress Δ ribosomes, New England Biolabs). Biotin was incorporated co-translationally at the two N-terminal amber positions TAG using the suppressor tRNA technique^4^. The system was supplemented with 10 μM of a modified tRNA pre-charged with biotinylated lysine (Biotin-XX-AF_tRNA, amber, CloverDirect), 0.5-5 μM trigger factor, as well as murine RNase inhibitor (New England Biolabs). Synthesis was initiated by mixing the system with the *E. coli* ribosomes coupled to beads, and the 5.5 nM linear DNA template. The reaction mixture was incubated at 37°C for 30 min. The resulting nascent chains remained attached to the ribosome due to the SecMstr arrest peptide at the C-terminus^5,6^. Following the transcription/translation reaction, the bead with their tethered and stalled ribosome nascent chain complexes (RNCs) were resuspended in TICO buffer (20 mM HEPES-KOH pH 7.6, 10 mM (Ac)_2_Mg, 30 mM AcNH_4_, 4 mM β-mercaptoethanol) at 4°C. The RNC-coupled beads were diluted in 300 μL TICO buffer prior to their usage in the optical tweezers. As oxygen scavenger the P2O system (3 units per ml pyranose oxidase, 90 units per ml catalase and 50 mM glucose, Sigma) was used.

Preparation of the FlAsH Dye

Stock solutions of 5 mM were prepared by dissolving FlAsH-EDT_2_ (Carbosynth) in DMSO (Thermo Scientific) and stored in an inert atmosphere at −20°C. For experimental use, the stock solutions were diluted in TICO buffer to a working concentration of 100 nM.

Optical tweezers assay and single-molecule data analysis

Correlated single-molecule force spectroscopy and multi-color confocal laser scanning spectroscopy measurements were carried out with the C-trap instrument (Lumicks, Amsterdam). This instrument features two optical traps formed by a high intensity- and polarization-stable single 1064 nm laser, which is split into two orthogonally polarized beams, one of which can be steered with a piezo mirror relative to the other. Two fluorescence excitation lasers (532 nm and 638 nm) allow for dual color confocal fluorescence, while the dedicated APDs assure single-photon sensitivity. Measurements were performed in a monolithic laminar flow cell with a stable passive pressure driven microfluidic system with 5 separate flow channels. Data was acquired at a rate of 50 kHz, decimated/averaged down to 500 Hz, and was analysed using custom scripts in Matlab and python. Calibration of the two orthogonally polarized traps was performed using the power spectrum method, where the power spectra obtained from the beads undergoing Brownian motion in the optical traps are fitted with a Lorentzian to obtain conversion parameters for displacements and forces in nm and pN^7^. The trapping laser intensity was kept constant for all measurements, resulting in a trap stiffness of about 260 ± 50 pN/μm. In order to tether individual molecules, the optically trapped beads were brought within close proximity for short time intervals, before separating them to an inter-bead distance of about 3 μm. A slight increase in the force would signal tether formation. Measurements were taken in a cycling force spectroscopy mode, where the steerable trap was moved at a constant rate of 0.2 μm/s between a minimum bead separation of 2 μm and a maximum force of up to 65 pN. The resulting force-extension curves for individual tethers were fitted with two worm-like chains (WLC) in series, one for the DNA handle contribution (extensible worm-like chain, eWLC)^8^ and the other for the stalled nascent chain contribution (inextensible worm-like chain, WLC)^9^, yielding an average DNA persistence length of 43 ± 10 nm (SD of 5 nm) and a DNA stretch modulus of 1037 ± 388 pN/nm (SD of 194 pN/nm). Tethered molecules undergoing only gradual transitions could not be fitted with the Odijk inextensible approximation WLC model, and hence WLC rulers were used corresponding to fully compacted and fully extended monomer and dimer chains as depicted in Fig. 3B, 4B, and Extended Data Fig. 3, 4, 7, 8, 9 using the average DNA parameters obtained above. Four features were identified during the pulling-relaxation experiments of monomer and dimers. Most of the total chain was characterized as: (i) unfolded, (ii) extending or compacting gradually, (iii) compacted and unfolding in discrete events, and (iv) compacted and not unfoldable. More specifically, the features were: (i) An unfolded state, in which more than half of the total monomer or dimer chain is unfolded, and remained so during a stretch-relax cycle. (ii) A compacted (or an extended) state composed of more than half of the total monomer or dimer chain, which displayed a gradual contour length increase (decrease) of more than half of the total chain length during a stretch-relax cycle. (iii) A compacted state composed of more than half of the total monomer or dimer chain, which displayed multiple discrete contour length changes totalling more than half of the total chain length during a stretch-relax cycle. (iv) A compacted state composed of more than half of the total monomer or dimer chain, which displayed no detectable contour length change during a stretch-relax cycle.
